# Consideration of species-specific diatom indicators of anthropogenic stress in the Great Lakes

**DOI:** 10.1101/514273

**Authors:** Euan D. Reavie, Meijun Cai

## Abstract

Robust inferences of environmental condition come from bioindicators that have strong relationships with stressors and are minimally confounded by extraneous environmental variables. These indicator properties are generally assumed for assemblage-based indicators such as diatom transfer functions that use species abundance data to infer environmental variables. However, failure of assemblage approaches necessitates the interpretation of individual dominant taxa when making environmental inferences. To determine whether diatom species from Laurentian Great Lakes sediment cores have the potential to provide unambiguous inferences of anthropogenic stress, we evaluated fossil diatom abundance against a suite of historical environmental gradients: human population, agriculture, mining, atmospheric nutrient deposition, atmospheric temperature and ice cover. Several diatom species, such as *Stephanodiscus parvus*, had reliable relationships with anthropogenic stress such as human population. However, many species had little or no indicator value or had confusing relationships with multiple environmental variables, suggesting one should be careful when using those species to infer stress in the Great Lakes. Recommendations for future approaches to refining diatom indicators are discussed, including accounting for the effects of broad species geographic distributions to minimize region-specific responses that can weaken indicator power.

## Introduction

In a report on 50 priority research questions in paleoecology [1], two of the priorities were as follows and are the subject of the current analysis: What methods can be used to develop more robust quantitative paleoenvironmental reconstructions? How can paleoecologists disentangle the separate and combined effects of multiple causal factors in paleoecological records?

Knowing indicator properties of species is critical for robust inferences of environmental condition in monitoring programs and paleolimnological applications. In figuring out these species characteristics, the present-day distributions of diatoms (or any of several indicator organisms) are typically calibrated across the gradient of a selected environmental parameter [2]. In Anthropocene studies these variables tend to be analytes related to stressors such as nutrient enrichment (e.g. phosphorus), acidification (e.g. pH) and warming (e.g. salinity). These tools are usually assemblage-based and use a species weighted-average approach and have consistently provided valuable tools to infer past conditions from sedimentary records [3]. Monitoring programs often use biological indices such as the diatom-based Trophic Diatom Index (TDI) [4], which assigns a single environmental condition based on a weighted compilation of species characteristics for all taxa within a sample. Such distilled indices are desirable for aquatic management, and there is little question that strong relationships often exist between the abundance and structure of diatom communities and prevalent stressor variables. However, the limitations of aggregate community metrics were recognized by King & Baker [5], who emphasized that it is critically important to know which taxa are affected by a stressor of interest. This is especially important when aggregate metrics fail, such as recently occurred when applying a phosphorus transfer function to fossil diatom assemblages in Lake Huron [6].

When we examine modern diatom assemblages and fossil profiles, there is a desire to have clearly-defined stressor responses for our indicator species so that we may make clear interpretations of past conditions. This is challenging for many species due to their simultaneous responses to many environmental variables. Take *Fragilaria crotonensis* as an example. This pennate, phytoplanktonic diatom is common in the Laurentian Great Lakes and tends be most common during summer months [7]. Based on Reavie et al. [7], this taxon has relatively high optimum for phosphorus and turbidity and a low optimum for nitrogen. It has been noted in paleoecological studies as increasing in relative abundance following artificial phosphorus enrichment resulting in cultural eutrophication [8]. In an experimental study, *F. crotonensis* increased in response to nitrogen (and not silica or phosphorus) enrichment [9], an observation backed by observations of natural assemblages from Lake Michigan [10]. It is apparently also positively supported by water column mixing that enables heavy colonies to remain suspended [11]. Clearly, the multi-variable context of an aquatic environment has a critical bearing on the abundance of *F. crotonensis* and what it indicates relative to anthropogenic stressors. Such dynamics make indicator interpretations challenging, and furthermore, uncertainties in explaining the presence of a species is subject to unexplained noise in the data, which is expected when one deals with highly complex biological data such as diatom assemblages. These complicated indicator relationships likely contribute to poor performance of diatom-based indicator models, something that has been highlighted relatively recently [12].

A useful index requires accurate environmental calibration of species. However, geographic constraints can limit the large sample size needed to adequately calibrate diatom indicators [13]. Training sets of samples used to calibrate diatom indicators are preferably selected within a narrow range of physicochemical conditions that at the same time maximize the gradient of the stressor variable. Such an ideal dataset is rarely possible, so it is typical to include deep and shallow lakes that, respectively, may favor phytoplanktonic or benthic species, which is fine if one hopes to reconstruct changing lake depth and not some chemical parameter. Variations in catchment soil and bedrock determine ionic properties of aquatic systems, which in turn naturally drive assemblage properties, a phenomenon that determines diatom assemblages throughout the Laurentian Great Lakes basin [7]. Hence, the natural context of water quality can limit how a species responds to stress, if it even occurs at all, under certain natural conditions. Such geographic considerations are likely a major confounding factor in diatom indicator development, yet to our knowledge corrections for geographic context have not been applied in diatom-based transfer functions.

Aggregate metrics such as transfer functions and biological indices will continue to be undermined by multivariate determinants of diatom occurrence, so it is likely that species-specific interpretations will continue to be required for useful inferences of condition. We aimed to determine whether diatom species exist that have clear relationships with stress. Investigation was based on a suite of fossil diatom data and historical stressor data on population growth, agriculture, deforestation and climate change in the Laurentian Great Lakes Basin. Consideration is given to the importance of spatial variation and whether it can be overcome in the pursuit of indicator development.

## Methods

### Diatom dataset

Sediment cores were collected from 10 locations throughout the Great Lakes, as detailed by Reavie et al. [14]. Cores were age dated using ^210^Pb, and details of those age models are provided by Reavie et al. [15] (in review). Diatom frustules in sediment samples were cleaned and prepared for light microscopy. Diatoms were identified and enumerated at 1000 - 1250 X magnification with oil immersion [16].

For every sediment sample analyzed for a diatom assemblage, we generated associated stressor data for human population, agriculture, forest cover and atmospheric variables (annual minimum temperature, water level, NH_4_^+^, NO_3_^−^, inorganic N and chloride). Historical data for a given sample were selected based on the ^210^Pb-inferred date in the middle of the sample interval. The long-term stressor data (Supplement A) were compiled several ways:
 
- Quantitative data for agriculture, forests and population were retrieved from a collection of historical data compiled by Reavie et al. [17]. These data were compiled from various Canadian and US historical records and extrapolated, if necessary, into yearly estimates of stressor values going back to 1780. This stressor database is organized according to 60 watersheds comprising the Great Lakes watershed. Stressors for a given core location were associated according to adjacent sub-watersheds as depicted in Fig 1.
- Historical monthly minimum, maximum and mean air temperature data from mid 1800s to present were summarized for each of five Great Lakes by averaging the monthly minimum, maximum and mean data of meteorological stations assigned to each lake. Meteorological stations relevant to each lake were selected by Hunter et al. [18]. The monthly minimum, maximum and mean air temperature of each station was computed from daily data. Daily air temperature data were downloaded from the National Climatic Data Center (NCDC) [19] of the National Oceanic and Atmospheric Administration (NOAA) and from Canadian Weather [20]. The monthly minimum and maximum temperatures were calculated by taking the smallest and largest daily temperatures within that month. Because NCDC weather data did not give daily mean temperatures, the daily mean temperature was calculated by averaging daily minimum and maximum temperatures. Although we compiled these variations of temperature data, we ultimately selected analyses using annual minimum temperature due to its apparent dominant effect on phytoplankton communities in the Great Lakes [14] and because, worldwide, it is the most rapidly increasing temperature parameter [21].
- The atmospheric deposition monitoring stations within 100 km distance from a given lake’s shoreline were selected to calculate loading relevant to that lake. Annual wet deposition loadings of ammonium, nitrate, inorganic nitrogen and chloride were downloaded from National Atmospheric Deposition Program (NADP) [22] and EPA Clean Air Status and Trends Network (CASTNET) [23] stations from 1979 to 2014. A total of 52 monitoring stations (34 currently active stations and 18 historical stations) were relevant to the 100-km Great Lakes buffer. Some stations were within 100 km of more than one lake. The loadings for a lake were calculated by using the following distance weighted-average formula: 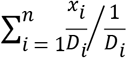, where *n* is the number of stations within 100 km of the lake, *x*_*i*_ is the loading at station *i* and *D*_*i*_ is the distance of station *i* from the lake shoreline.
- Monthly ice cover concentration and annual ice duration of winter (December to May) maps from 1973 to 2013 were obtained from GLAHF [24]. GIS maps were computed in ArcMap using the zonation tool to get mean monthly ice cover area and annual duration for each lake. The monthly ice cover from December (of the previous year) through May were averaged to get mean ice cover.
- Annual average water levels from 1860 to 2014 were downloaded from EPA [25].

**Fig 1.**
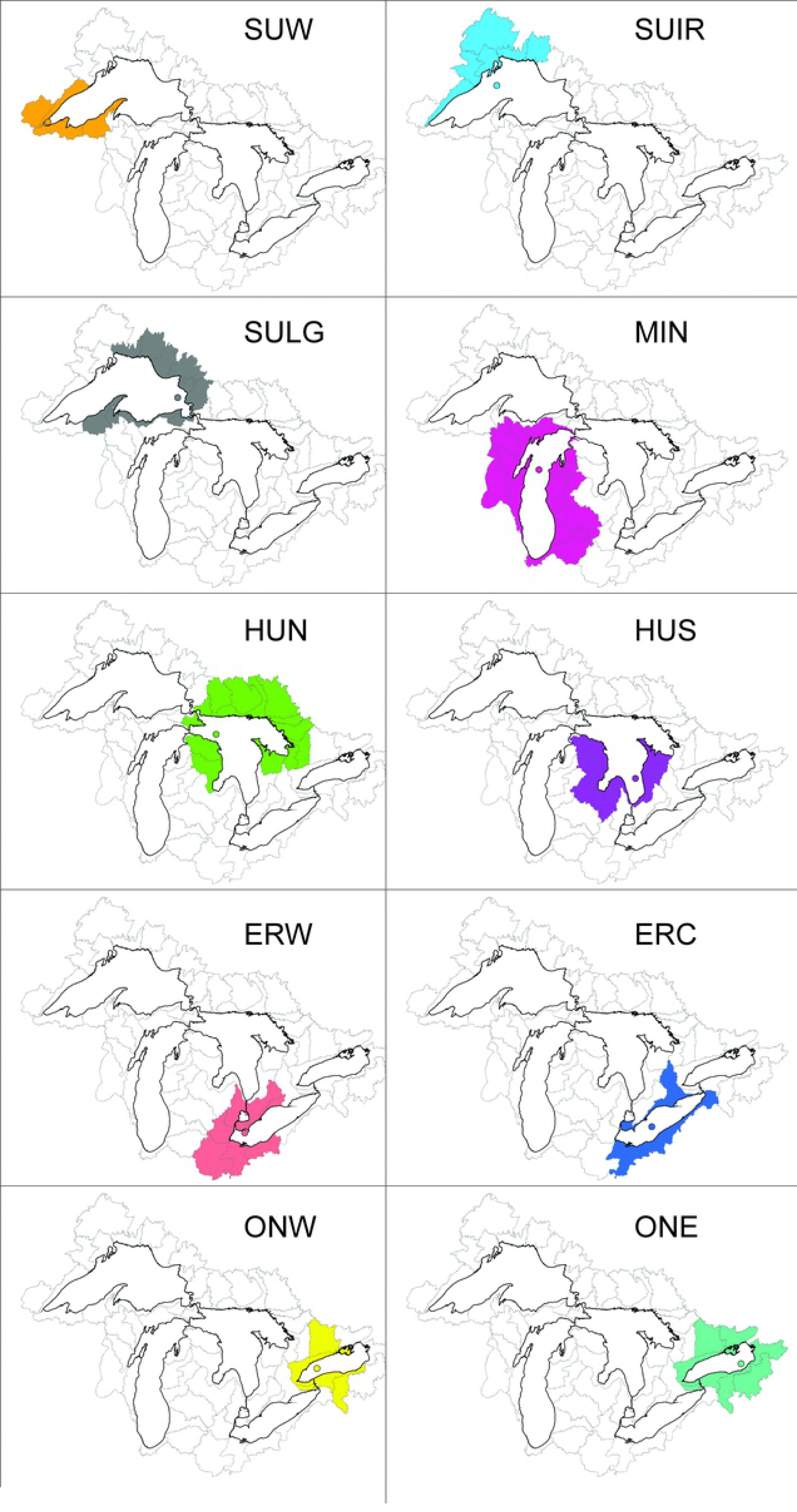
Maps of the Laurentian Great Lakes catchment, indicating the sub-watersheds selected to relate stressors with the sedimentary record in each sediment core (circle symbols) Delineation of sub-watersheds and quantitative stressor data are available from Reavie et al. (2018).

To evaluate species-stressor relationships, sample relative abundances of individual diatom species (at least three non-zero occurrences) were related to historical stressor data using Pearson correlation. This was first done using raw stressor data; then, for evaluations against temperature, atmospheric deposition and water level data, the analysis was performed again using standardized data. Standardization included transforming the stressor (x axis) and diatom relative abundance (y axis) data from a specific lake so that their standard deviations ranged from 0 to 1. After lake-specific standardization, the data were re-pooled for analysis. Standardization was deemed necessary due to major geographic differences in assemblage and stressor data. For instance, Lake Superior contains high relative abundances of cyclotelloid centric diatoms, more so in recent years, apparently due to recent warming [21]. However, because Lake Superior has always had high abundances of cyclotelloids, and because it is the coldest of the Great Lakes, regressions using raw data suggested that colder atmospheric temperatures favored higher relative abundances of *Cyclotella*. This was contrary to findings that increasing temperatures in a given lake tend to favor cyclotelloids. Hence, standardization by lake allowed correlation analyses to account for historical changes noted in paleolimnological records. The debatable appropriateness of this standardization is considered further in the Discussion section.

## Results

Correlations with stressor variables identified several species likely to be good indicators of stress. For the strictly anthropogenic stressors (population, agriculture, deforestation, mining), *Stephanodiscus parvus* (STEPARV) is ranked the highest, having the strongest relationship (highest r and lowest P) with higher human populations around the Great Lakes (Table 1). This is followed by *Stephanodiscus alpinus*, *Aulacoseira granulata* and *Fragilaria crotonensis*, all of which were strongly and positively correlated with human population. Due to its relative dominance in the earliest years represented in our sediment cores, *Cyclotella ocellata* had large, negative relationships with population and agriculture and a positive relationship with forestry. A total of 151 significant (with Bonferroni correction) species-stressor relationships occurred, 39 for agriculture, 26 for deforestation, 22 for mining stress, 28 for new mines and 36 for population. Using an uncorrected P (0.05), 668 significant relationships occurred. 83.6% (25.7% with Bonferroni correction) of the 335 taxa with sufficient occurrences for analysis had significant relationships with at least one stressor.

**Table 1.** Pearson correlations (r) of diatom relative abundance versus anthropogenic stressors. The top 25 taxon-stressor relationships are presented, sorted by smallest P value. N = number of observations in samples. A full list of the significant taxon-stressor relationships is in Supplement B.

| Diatom code | Full name | Variable | r | P | N |
| --- | --- | --- | --- | --- | --- |
| STEPARV | *Stephanodiscus parvus* Stoerm. & Hak. | Population | 0.70 | 0.0E+00 | 252 |
| STEALPI | *Stephanodiscus alpinus* Hust. | Population | 0.65 | 0.0E+00 | 252 |
| AULGRAN | *Aulacoseira granulata* (Ehr.) Simonsen | Population | 0.61 | 0.0E+00 | 252 |
| FRACROT | *Fragilaria crotonensis* Kitton | Population | 0.57 | 0.0E+00 | 252 |
| TABFENE | *Tabellaria fenestrata* (Lyngb.) Kütz. | Mining | 0.53 | 0.0E+00 | 254 |
| CYCOCEL | *Lindavia ocellata* (Pant.) Nakov, Guillory, M.L.Julius, E.C.Ther. and A.J.Alverson | Mining | 0.49 | 0.0E+00 | 254 |
| TABFENE | *Tabellaria fenestrata* (Lyngb.) Kütz. | New Mines | 0.49 | 0.0E+00 | 254 |
| FRACAPR | *Fragilaria capucina* var. *rumpens* (Kütz.) Lange-Bertalot ex Bukht. | New Mines | 0.49 | 0.0E+00 | 254 |
| CYCPSEU | *Discostella pseudostelligera* (Hust.) Houk & Klee | Population | -0.49 | 0.0E+00 | 252 |
| STEPSBTR | *Stephanodiscus subtransylvanicus* Gasse | Agriculture | -0.52 | 0.0E+00 | 249 |
| CYCOCEL | *Lindavia ocellata* (Pant.) Nakov, Guillory, M.L.Julius, E.C.Ther. and A.J.Alverson | Agriculture | -0.53 | 0.0E+00 | 249 |
| CYCOCEL | *Lindavia ocellata* (Pant.) Nakov, Guillory, M.L.Julius, E.C.Ther. and A.J.Alverson | Population | -0.53 | 0.0E+00 | 252 |
| CYCCOMRC | *Lindavia delicatula* (Hust.) Nakov, Guillory, M.L.Julius, E.C.Ther. & A.J.Alverson | Agriculture | -0.53 | 0.0E+00 | 249 |
| ACHMINUS | *Achnanthes minutissima* var. *scotica* (Carter) Lange-Bertalot | Agriculture | -0.56 | 0.0E+00 | 249 |
| CYCPSEU | *Discostella pseudostelligera* (Hust.) Houk & Klee | Agriculture | -0.56 | 0.0E+00 | 249 |
| SYNOSTE | *Synedra ostenfeldii* (Krieg.) A. Cl. | Agriculture | -0.57 | 0.0E+00 | 249 |
| SYNFILIE | *Synedra filiformis* var. *exilis* A. Cl. | Agriculture | -0.58 | 0.0E+00 | 249 |
| STEPARV | *Stephanodiscus parvus* Stoerm. & Hak. | Forestry | -0.48 | 4.4E-16 | 253 |
| RHIERIE | *Urosolenia eriensis* (H.L.Sm.) Round & R.M.Crawford | Agriculture | -0.48 | 4.4E-16 | 249 |
| CYCATOM | *Cyclotella atomus* Hust. | Agriculture | -0.48 | 8.9E-16 | 249 |
| RHIGRAC | *Rhizosolenia gracilis* H.L.Smith | Agriculture | -0.46 | 9.8E-15 | 249 |
| ENTSP | *Entomoneis* Ehrenb. spp. | New Mines | 0.46 | 1.0E-14 | 254 |
| FRABREV | *Pseudostaurosira brevistriata* (Grunow) D.M.Williams & Round | Population | 0.46 | 2.0E-14 | 252 |
| ACHMINUS | *Achnanthes minutissima* var. *scotica* (Carter) Lange-Bertalot | Population | -0.46 | 2.1E-14 | 252 |
| STEPSBTR | *Stephanodiscus subtransylvanicus* Gasse | Population | -0.46 | 2.7E-14 | 252 |

For the stressor variables with natural components (chemical deposition and atmospheric temperatures), *Stephanodiscus alpinus* type I had the highest, positive correlations with nitrate and inorganic N deposition (Table 2). The next highest ranked were *S. parvus* (positively related to minimum annual temperature) and seven species that were negatively related to minimum annual temperature (*Rhizosolenia eriensis*, *Rhizosolenia gracilis*, *Synedra filiformis* v. *exilis*, *Achnanthidium minutissimum*, *C. ocellata*, *Synedra ostenfeldii* and *Stephanodiscus subtransylvanica*). A total of 126 significant (with Bonferroni correction) relationships occurred, 19 for chloride, 21 for inorganic N, 17 for ammonium, 21 for nitrates and 48 for minimum annual temperature.

**Table 2.**
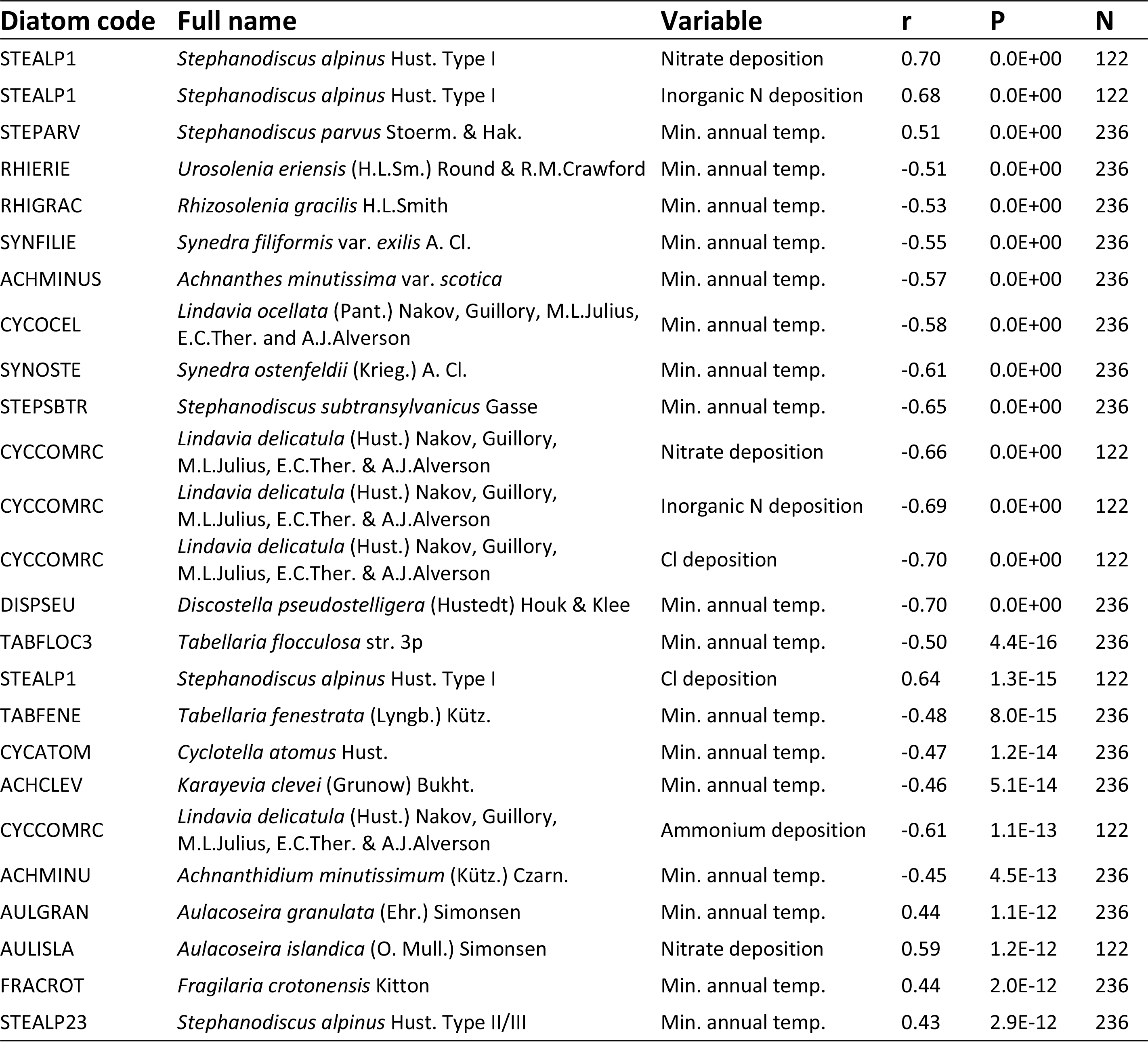
Pearson correlations (r) of diatom relative abundance versus measured environmental variables considered to represent a mixture of natural and anthropogenic components. The top 25 taxon-variable relationships are presented, sorted by smallest P value. N = number of observations in samples. A full list of the significant taxon-stressor relationships is in Supplement B.

Lake-based normalization of fossil and historical monitoring data resulted in a shift in the indicator properties of several diatom species and resulted in fewer significant relationships with stressors (Table 3), indicating geographical specifics can influence interpretations of condition. With Bonferroni correction applied, only 11 taxa had significant correlations, all with minimum annual temperature. As anticipated, many (5) of these relationships were positive correlations for taxa belonging to *Cyclotella sensu lato*. Examining the regressions on two select taxa (Fig 2) indicates the effect of within-lake data standardization. The relationship between *Lindavia laurentiana* and temperature became stronger after standardization, whereas the relationship for *Lindavia delicatula* reversed, becoming positive and corresponding with that observed in paleorecords (i.e. increasing in relative abundance with higher atmospheric temperatures) [21].

**Table 3.**
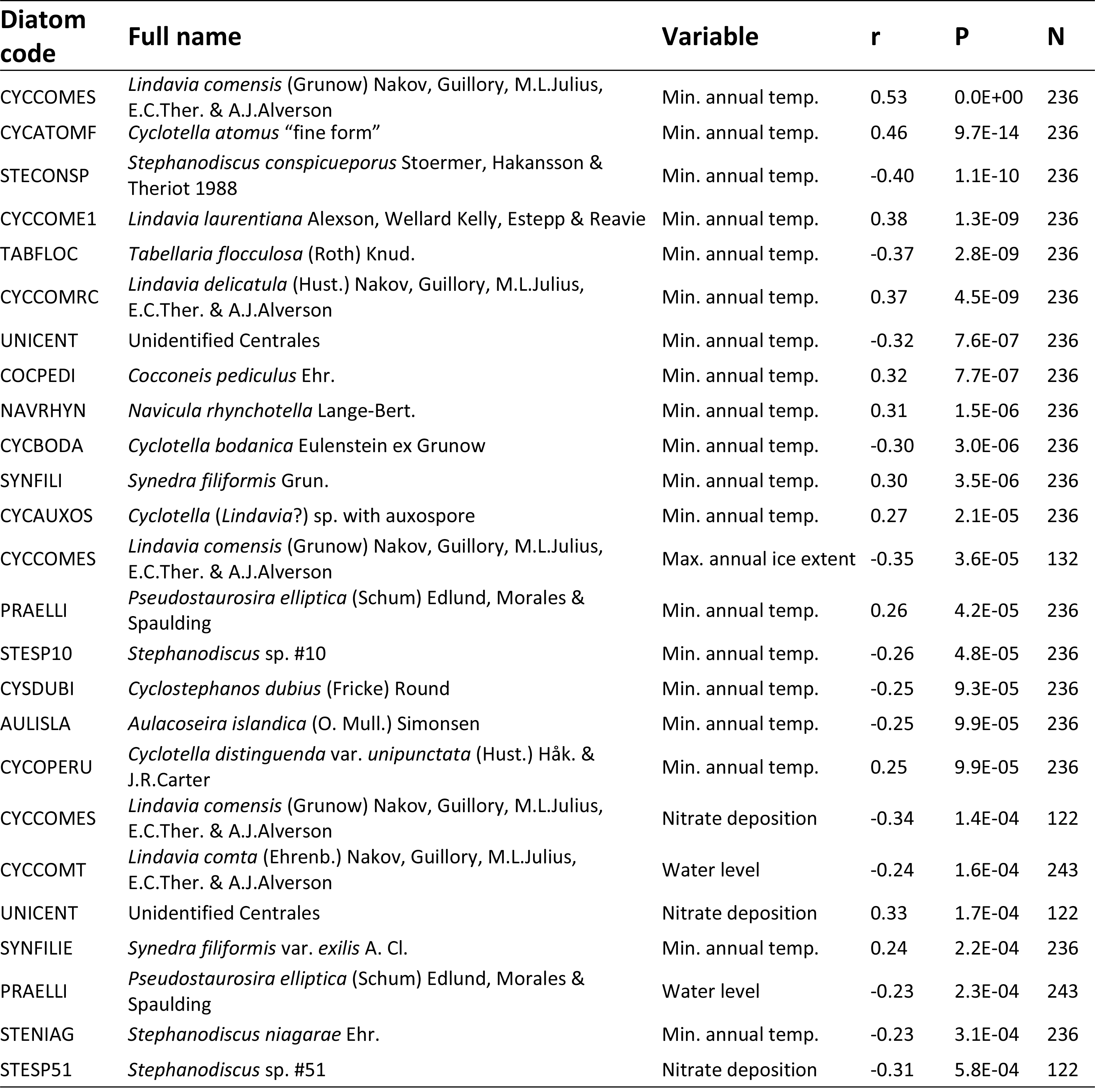
Pearson correlations of diatom relative abundance versus measured physical parameters and chemical atmospheric deposition. For this analysis diatom and environmental data were standardized by lake so that SDs ranged from 0 to 1. The top 25 taxa are presented, sorted by smallest P value. N = number of observations in samples. A full list of the significant taxon-stressor relationships is in Supplement B.

**Fig 2.**
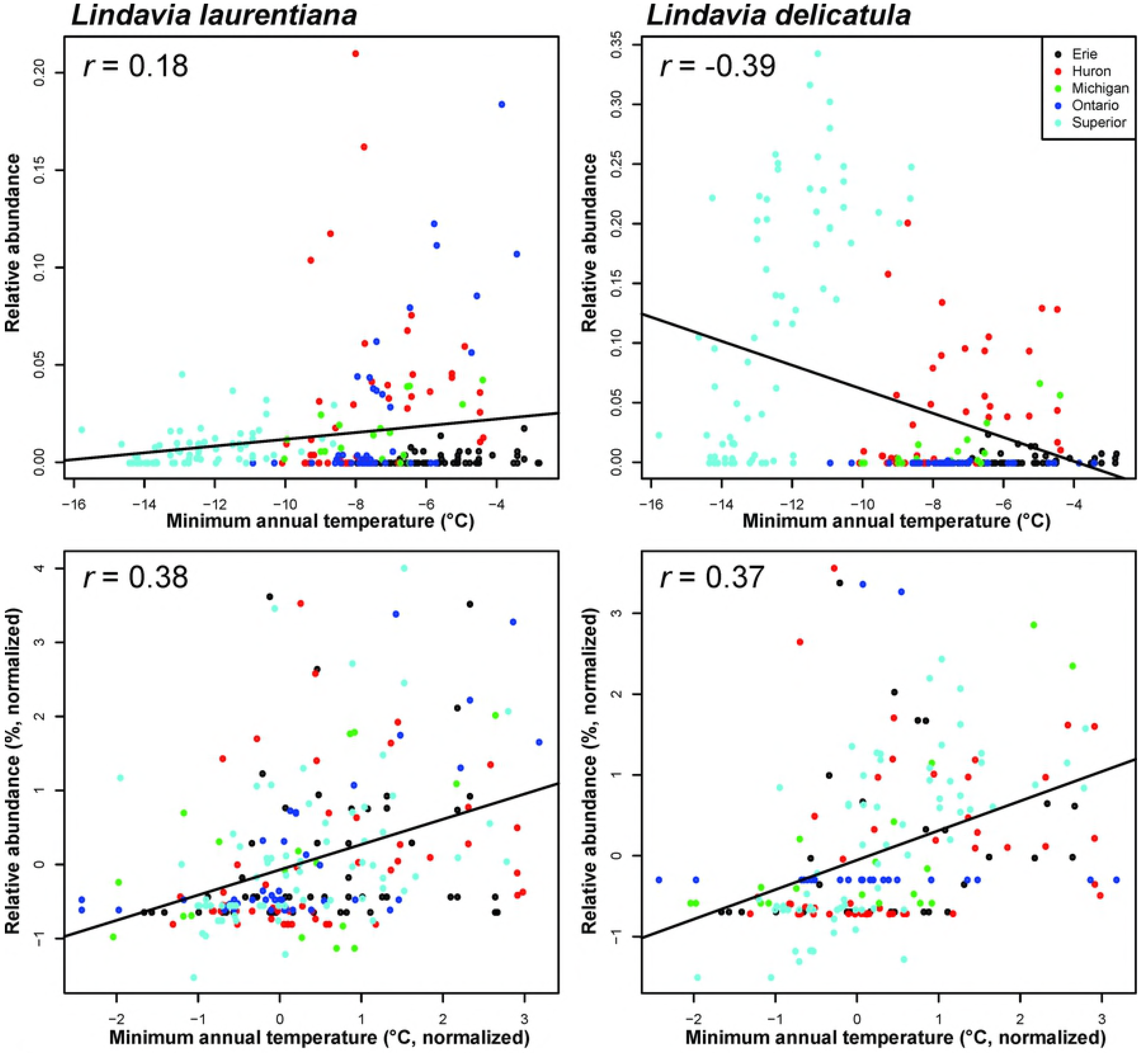
Temperature-diatom relationships for two cyclotelloid species in the Great Lakes. The upper plots use the original temperature and percent abundance data, and lower plots use data that have been standardized by lake and recombined.

## Discussion

We identified several diatom taxa that individually reflect stress, or lack of stress, in the Great Lakes. In general, taxon-stressor relationships with the lower P and higher r values (Tables 1-3) can be considered better indicators. While there are too many taxa to discuss, high-ranking species like *S. parvus* should be considered good indicators of stress associated with population growth, agriculture and deforestation in the catchment of an aquatic system. Such a finding is not necessarily novel for a high-nutrient indicator such as *S. parvus*, but our results suggest this indicator status is minimally confounded by environmental variability throughout the Great Lakes. As one steps down the ranked lists of species, greater caution should be applied when associating a taxon with a particular stressor.

Diatoms have existed for millions of years, well before freshwater assemblages began to shift due to human drivers. Hence, unlike synthetic fossil indicators (such as PCBs), a diatom species cannot explicitly indicate anthropogenic stress. However, it is possible that pre-impact paleorecords may be used to strengthen species indicator characteristics. For instance, did *S. parvus* exist in high abundance prior to the Anthropocene? A lack of such a record would strengthen the characterization of *S. parvus* as an indicator of stress. In Great Lakes containing *S. parvus* in modern records, the relative abundance of *S. parvus* was consistently low in fossil records prior to the 20^th^ century [26,27]. While not a complete assessment of the paleolimnology literature, a review of several manuscripts where the species was observed [28–31] indicates that *S. parvus* was not present in significant numbers prior to human population expansion and associated nutrient enrichment of aquatic systems. Even in a Swiss lake with a long record of human impacts, *S. parvus* did not achieve notable abundance until catchment pasturing as early as the 14^th^ century [32]. Indeed, all of the Great Lakes now contain dominant species that were not previously observed, or existed in trace amounts, prior to the Anthropocene. Several species in the *Cyclotella sensu lato* complex, especially *Lindavia delicatula* and *Lindavia comensis*, are clearly dominant due to human changes to Great Lakes conditions [21]. While outside the scope of this paper, such scanning of available literature may be a worthwhile method to qualify the indicator properties of species.

Although we provide some clarifications, confirming an independent response of a species to stress remains challenging. A species’ consistent response to stressors such as population, agriculture and deforestation is helpful, but most taxa reveal relationships with variables with counterintuitive meaning. For example, based on correlations, *Achnanthidium minutissimum* is associated with a mixture of stressors: low temperatures, agriculture, population and nitrogen deposition, and seemingly in conflict, high mining influence. Bradshaw and Anderson [33] noted that the performance of a diatom-based transfer function for phosphorus was strongly driven by one species, *Cyclostephanos dubius*, which has a high phosphorus optimum. Subsequent analyses identified a strong relationship between *C. dubius* abundance and nitrate in five Danish lakes. In a case like this, inferences of productivity may be sufficient since these two variables tend to co-vary. But what if the variables were phosphorus and pH? Such a finding would complicate interpretations of condition because those variables typically indicate different stress types. Saros [34] summarizes several examples of experiments indicating the confounding effects of multivariate control over species abundance has on the responses of those species to an environmental variable one may wish to reconstruct. Further analysis of the independence of species relationships with a given stressor may help further refine these indicators, but a more detailed set of measured variables than available to us would be needed (e.g. long-term records of phosphorus, conductivity, grazer and filter feeder abundance, water clarity).

Where do we go from here? As recommended by Saros [34], additional indicators (whether or not they employ diatoms) should be considered. We have focused on diatom taxonomic composition of samples, but we reiterate what other researchers have said [35]: several possible complementary indicators exist to provide a more refined or diverse interpretation of condition from paleorecords. Additional biological (e.g. chironomids, chrysophytes), chemical (e.g. geochemistry, isotopes) and physical (e.g. grain sizes) indicators are at our disposal, should funds allow.

To expand the utility of these indicators beyond the Great Lakes system may require input from experts, a method that had success on a diatom-based indicator of stream condition in the northeast US [36]. Briefly, this approach employed diatom experts to vote on the indicator properties of taxa based on available data and their experience as diatom researchers. While it is difficult to quantify such associations between diatom species and stress, expert input is likely to apply historical knowledge from outside the immediate study area and so is likely to provide a broader context on a diatom species’ indicator status.

Could indices and transfer functions be improved by eliminating mundane or highly confounded species? This has been explored by Racca et al. [37] on a set of arctic lakes using a neural network approach that pruned species from a training set according to their unique influence by the variable of interest. This appeared to improve transfer function performance. Unfortunately, since publication of that study we can find no similar applications, though other methods for removing non-responsive taxa [38,39] and sample locations having extreme values of a confounding variable (e.g. elevation) [40] from transfer functions appear to be promising. Perhaps future applications should consider a permanent method for determination of confounded species prior to transfer function use.

Isolating species cultures and evaluating against stressor surrogates (e.g. mesocosm or “bottle” studies of phosphorus concentrations) have been applied to better understand species responses (e.g. Saros et al.) [41]. Such studies are impractical for the thousands of diatom taxa that have been observed in modern and fossil records in the Great Lakes, but assessments of the most common taxa could be helpful. Further, it has long been recognized that species act differently under culture due to the confining of environmental condition that prevents accurate characterization of in-situ species and assemblage behavior [42].

If we assume that pre-standardization of data by geographic location is appropriate, then it follows that developing indicator tools such as transfer functions that rely on water quality will be negatively affected if geographic correction is not applied. Clearly, a species like *Lindavia delicatula* (Figure 2) will have very different indicator coefficients using data with and without spatial standardization. Further, it is highly likely such geographic artefacts have had important negative influences on transfer functions, hence the recommendation to ensure narrow environmental variation outside the reconstructed variable of interest [43]. Certainly, applying a transfer function using data calibrated using all of the Great Lakes [7] should be performed carefully, employing validation that fossil assemblages indicate assemblage responses to the variable of interest. Geography is one consideration, but other confounding variables include water quality variables that may not be measured during the species calibration phase.

This study has several outcomes: (1) There are several diatom taxa that are strong indicators of environmental stress and should be citable as such; (2) Many (most?) diatom taxa are apparently poor indicators of these stressors due to mundane responses or confounding by multiple variables, so caution should be used when making inferences based on these species; and (3) Geographic considerations are extremely important when calibrating diatom indicators. While constraining a training set geographically can reduce sample size, there are methods such as spatial normalization that may be useful to account for these confounding spatial factors.

## Acknowledgements

We thank K. Kennedy, A.R. Kireta, R.W. Sterner, C.A. Stow and the Research Vessel Lake Guardian and Blue Heron field crews for their help collecting core samples. Sediment dating was supported by D.R. Engstrom and personnel at the St. Croix Watershed Research Station. We thank Sonja Hausmann for input on a previous draft of this manuscript.

